# Structural basis for recruitment of DAPK1 to the KLHL20 E3 ligase

**DOI:** 10.1101/414250

**Authors:** Zhuoyao Chen, Sarah Picaud, Panagis Filippakopoulos, Vincenzo D’Angiolella, Alex N. Bullock

**Author notes:** Correspondence (A.N.B.).

## Abstract

BTB-Kelch proteins form the largest subfamily of Cullin-RING E3 ligases, yet their substrate complexes are mapped and structurally characterized only for KEAP1 and KLHL3. KLHL20 is a related CUL3-dependent ubiquitin ligase linked to autophagy, cancer and Alzheimer’s disease that promotes the ubiquitination and degradation of substrates including DAPK1, PML and ULK1. We identified a ‘LPDLV’-containing recruitment site in the DAPK1 death domain and determined the 1.1 Å crystal structure of a KLHL20-DAPK1 complex. DAPK1 binds to KLHL20 as a loose helical turn that inserts deeply into the central pocket of the Kelch domain to contact all six blades of the β-propeller. Here, KLHL20 forms a salt bridge as well as hydrophobic interactions that include a tryptophan and cysteine residue ideally positioned for covalent inhibitor development. The structure highlights the diverse binding modes of circular substrate pockets versus linear grooves and suggests a novel E3 ligase for protac-based drug design.

## INTRODUCTION

Ubiquitination by E1-E2-E3 enzyme cascades is the major mechanism by which cells mark proteins for degradation, but can also facilitate protein trafficking, transcriptional control and cell signaling (Hershko and Ciechanover, 1998; Rape, 2018). The substrate specificity of protein ubiquitination is determined at the level of E3 ubiquitin ligases, which recruit their cognate substrate proteins to complete the enzyme cascade (Buetow and Huang, 2016). Kelch-like protein 20 (KLHL20, also known as KLEIP) is a member of the BTB-Kelch family that assembles with CUL3 and RBX1 to form a multi-subunit Cullin-RING E3 ligase (Geyer et al., 2003; Hara et al., 2004; Pintard et al., 2004). These complex E3 ligases use the RBX1 subunit to engage a charged E2-ubiquitin pair before transferring the ubiquitin to substrates captured by the BTB-Kelch protein (Genschik et al., 2013; Petroski and Deshaies, 2005). Catalysis occurs via CUL3 neddylation, which stabilizes the correct geometry of the complex for ubiquitin transfer (Duda et al., 2008).

Like other BTB-Kelch family members, KLHL20 utilizes multiple functional domains. The BTB and 3-box domains confer binding to CUL3, whereas the Kelch β-propeller domain serves as the substrate recognition domain (Canning et al., 2013; Lee et al., 2010). To date, the majority of substrates identified for KLHL20 are targeted for proteasomal degradation, suggesting their modification by Lys48-linked polyubiquitin chains (Chen et al., 2016). These include the substrates DAPK1 (Lee et al., 2010), PML (Yuan et al., 2011), PDZ-RhoGEF (Lin et al., 2011) and ULK1 (Liu et al., 2016). However, KLHL20 also plays an important role in protein trafficking by targeting coronin 7 to the trans-golgi network through atypical K33-linked polyubiquitination (Yuan et al., 2014).

The substrates of KLHL20 reflect its function in cellular stress responses, as well as its linkage to human disease (Chen et al., 2016). Transcription of the *KLHL20* gene is upregulated by the hypoxia-inducible factor HIF-1α leading to its overexpression in hypoxic tumor cells (Yuan et al., 2011). In this context, KLHL20 can promote tumorigenesis by degrading the tumor suppressor proteins DAPK1 and PML. In human prostate cancer patients, higher levels of KLHL20 (and low PML) were found to correlate specifically with high grade tumours (Yuan et al., 2011). Moreover, KLHL20 depletion in PC3 prostate cancer cells restricted the growth of tumor xenografts suggesting KLHL20 as a potential therapeutic target (Yuan et al., 2011). KLHL20 also plays a critical role in autophagy termination by degrading the pool of activated ULK1 (Liu et al., 2016). Thus, KLHL20 can restrict both apoptotic and autophagic cancer cell death. Importantly, interferon stimulation causes the sequestration of KLHL20 in PML nuclear bodies (Lee et al., 2010). This inhibitory mechanism allows DAPK1 to evade degradation and to accumulate to mediate interferon-induced cell death (Lee et al., 2010). Notably, the stress responses of KLHL20 also appear linked to neurodegeneration with KLHL20 RNA transcript levels being among the top 20 biomarkers for Alzheimer’s disease progression (Arefin et al., 2012; Gomez Ravetti et al., 2010).

Despite the growing number of substrate proteins identified for the 50 members of the BTB-Kelch family, there remains limited knowledge of their specific binding epitopes and consequently a lack of structural information for the corresponding E3-substrate complexes. Here, we investigated the binding of KLHL20 to DAPK1, which was the first reported substrate for this E3 ligase (Lee et al., 2010). Yeast two-hybrid studies previously mapped the interaction to the death domain of DAPK1 and the Kelch domain of KLHL20. Through a peptide scanning approach we identified a ‘LPDLV’-containing recruitment site within this DAPK1 region that bound to KLHL20 with low micromolar affinity. We also determined the crystal structure of their complex at 1.1 Å resolution revealing a distinct peptide binding mode compared to the previously determined structural complexes of KEAP1 and KLHL3 (Lo et al., 2006; Schumacher et al., 2014). The novel structure further identifies a hydrophobic substrate pocket that may be more attractive for small molecule inhibitor development than the highly charged surface of KEAP1.

## RESULTS

### Mapping of the DAPK1 binding motif for KLHL20 recruitment

The recombinant death domain of DAPK1 (Figure 1A) has been shown to display intrinsic disorder and a high propensity for aggregation making it unsuitable for structural studies (Dioletis et al., 2013). Given the lack of structural order, we set out to map the DAPK1 binding epitope using the SPOT peptide technology. We synthesized a peptide array to span the length of the DAPK1 death domain using 15-mer peptides and a three amino acid frameshift at each position. Probing of the array with recombinant 6xHis-KLHL20 Kelch domain and anti-His-antibody for detection revealed protein capture at two sites encompassing DAPK1 residues 1327-1350 and 1378-1395, respectively (Figure 1B). A control experiment indicated that the binding epitope was likely to reside within the N-terminal region since peptides from the second site also bound to the anti-His antibody alone marking them as likely false positives (Figure 1B).

**Figure 1.**
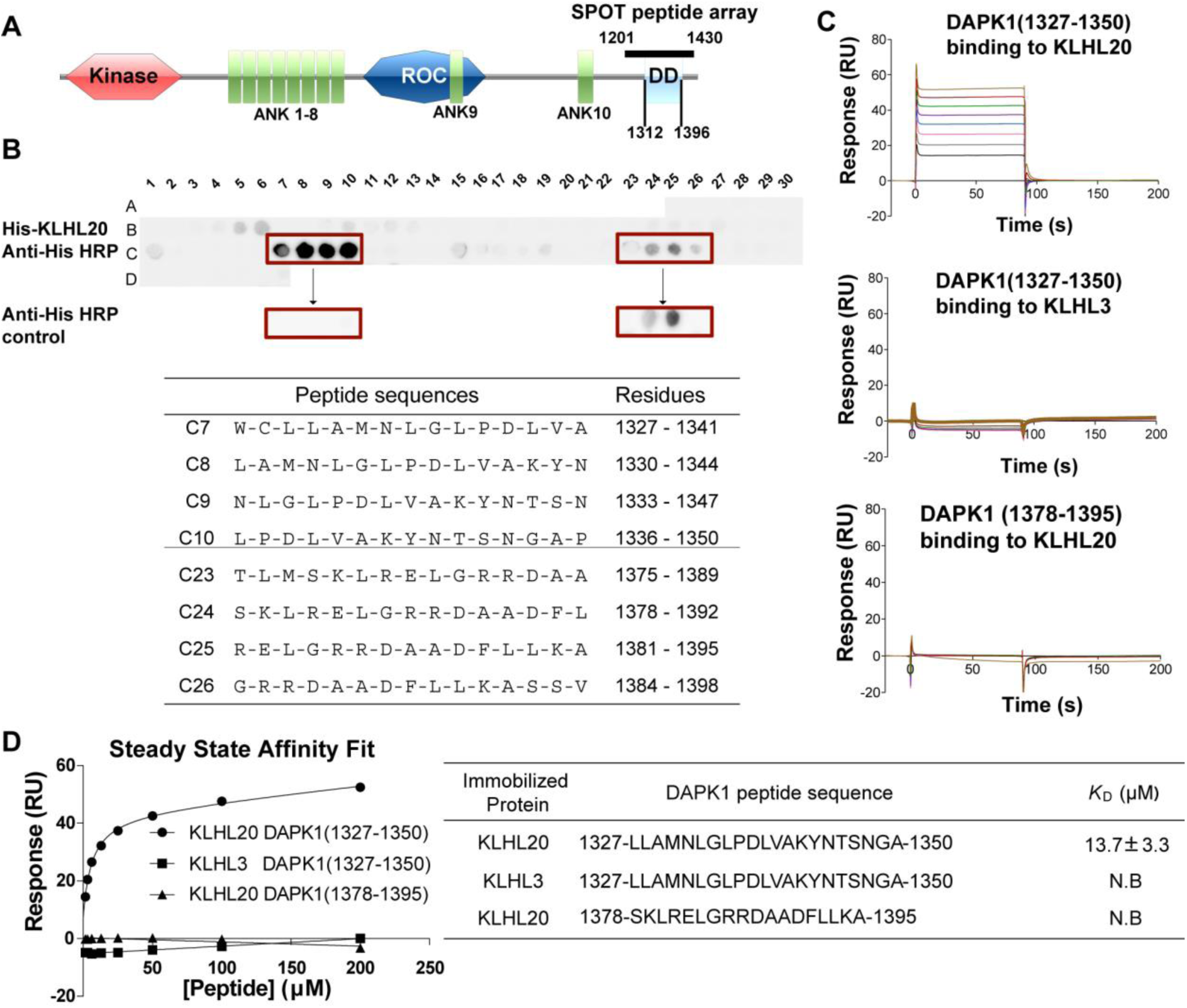
Mapping of the DAPK1 binding motif for KLHL20 recruitment. **(A)** Domain organization of human DAPK1 (ank, ankyrin repeat; DD, death domain comprising residues 1312 to 1396). Solid bar denotes the extended region explored for KLHL20 interaction (DAPK1 residues 1201-1430). **(B)** SPOT peptide array. Each spot was printed as a 15-mer DAPK1 peptide with a 3 residue frameshift at each consecutive position. Arrays were incubated with purified 6xHis-KLHL20 Kelch domain, washed and then KLHL20 binding detected using anti-His HRP-conjugated antibody. Binding was observed at two sites spanning DAPK1 residues 1327-1350 and 1378-1395, respectively. As a control, duplicate spots were probed with antibody alone and revealed nonspecific antibody binding to DAPK1 residues 1378-1395. **(C)** For SPR experiments, KLHL20 and KLHL3 Kelch domains were immobilized by amine coupling on different flow cells of a CM5 sensor chip. Indicated DAPK1 peptides were injected subsequently at concentrations of 1.6 μM, 3.1 μM, 6.2 μM, 12.5 μM, 25 μM, 50 μM, 100 μM and 200 μM. Binding was monitored at a flow rate of 30 μL/min. (**D**) SPR binding data were fitted using a steady state affinity equation. DAPK1 residues 1327-1350 bound to KLHL20 Kelch domain with *K*_D_ = 13.7 μM. (N.B., no binding detected).

To validate these putative interaction sites, we designed peptides for the two DAPK1 regions and performed surface plasmon resonance (SPR) experiments to measure their respective binding affinities for KLHL20 (Figure 1C). A DAPK1 peptide spanning the N-terminal site residues 1329-1349 bound robustly to the Kelch domain of KLHL20 with *K*_D_ = 13.7 μM (Figure 1D). The same peptide showed no apparent binding to the Kelch domain of KLHL3 demonstrating that the interaction was specific to KLHL20 (Figures 1D). A DAPK1 peptide spanning the C-terminal site residues 1378-1395 also failed to bind to KLHL20 confirming that this downstream region was a false positive (Figure 1D). Together these data identified a single epitope within the death domain of DAPK1 that showed both potency and specificity for interaction with KLHL20.

### An ‘LPDLV’ motif in DAPK1 is critical for KLHL20 interaction

Attempts to crystallize KLHL20 either alone or in complex with the identified 21-mer peptide from DAPK1 produced only microcrystalline material yielding poor diffraction. Therefore, we sought to refine the minimal DAPK1 epitope by using the SPOT technology for peptide truncation experiments, as well as alanine scanning to probe the sequence determinants of binding. The results from these experiments were in excellent agreement and identified DAPK1 residues Leu1336 to Val1340 as critical for KLHL20 interaction (Figure 2). N-terminal deletion or mutation of Leu1136 drastically reduced KLHL20 binding, whereas C-terminal deletion or mutation of Val1340 abolished all detectable binding (Figure 2). Other deletions and mutations outside of this region appeared well tolerated mapping the critical binding region to a ‘1336-LPDLV-1340’ motif in DAPK1.

**Figure 2.**
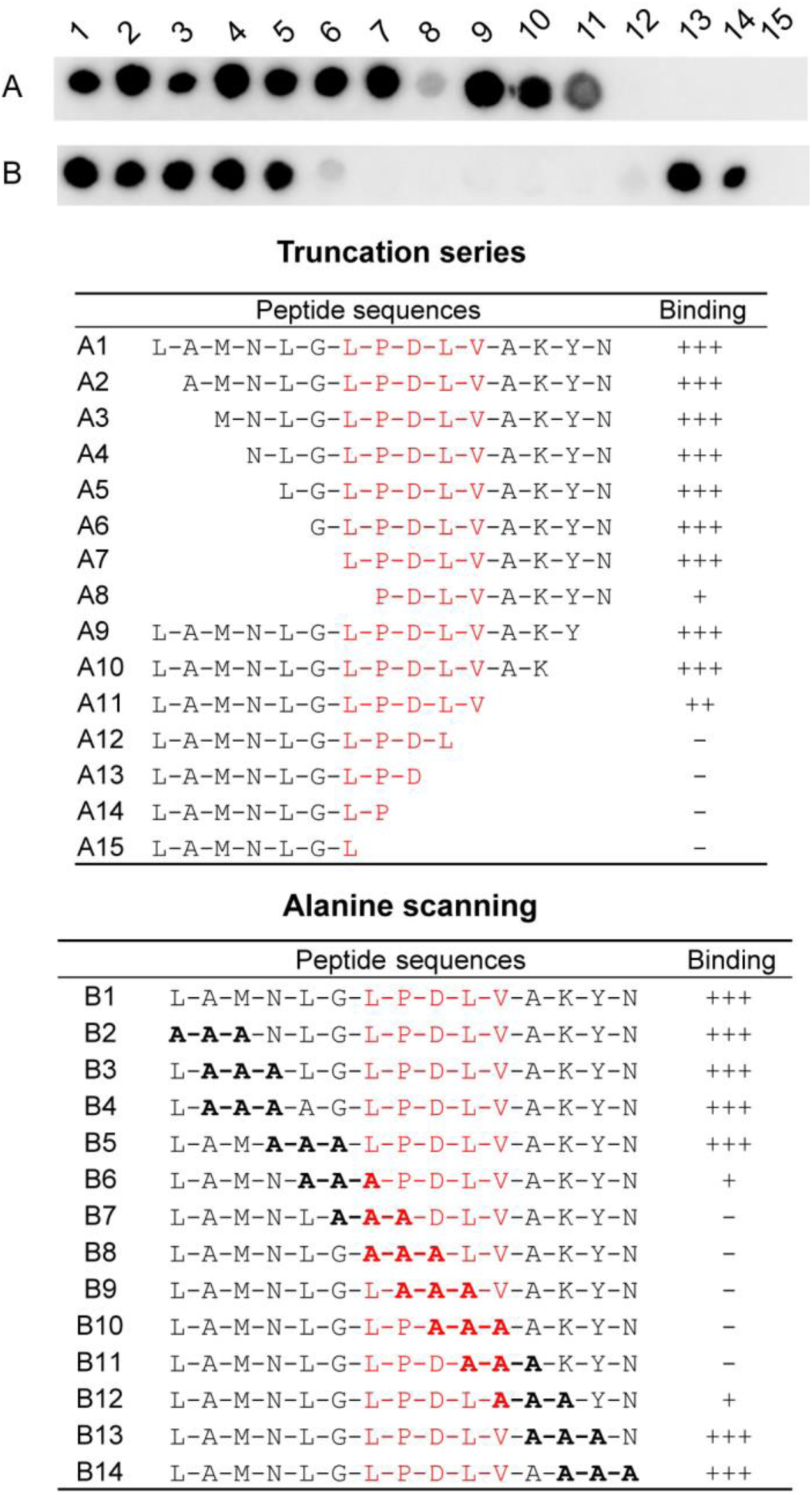
An ‘LPDLV’ motif in DAPK1 is critical for KLHL20 interaction. DAPK1 peptide variants were printed in SPOT peptide arrays. Row A peptides explored N and C-terminal truncations, whereas row B explored triple-alanine scanning mutagenesis. Arrays were incubated with purified 6xHis-KLHL20 Kelch domain, washed and then binding detected with anti-His antibody. KLHL20 binding was abrogated upon deletion or mutation of a central ‘LPDLV’ sequence motif in DAPK1.

### High resolution structure of KLHL20 bound to DAPK1 peptide

For further co-crystallization trials, we tried an 11-residue DAPK1 peptide (LGLPDLVAKYN) in order capture interactions of the central ‘LPDLV’ motif, while allowing for local conformational preferences and potential flanking interactions. Viable crystals were obtained with strong diffraction after a combination of microseeding from initial hits and fine matrix screening for further optimization. Subsequently, we were able to determine a high resolution structure for the complex of KLHL20 and DAPK1 peptide (Table 1). The structure was refined at 1.1 Å resolution and traced the full KLHL20 Kelch domain from residues 317 to 601 (Figure 3A). The complete DAPK1 peptide was also clearly defined in the electron density map (Figure 3B) allowing its binding interactions to be mapped in atomic detail.

**Table 1.** Data collection and refinement statistics for the KLHL20-DAPK1 complex.

| **Date collection** |  |
| --- | --- |
| Wavelength (Å) | 0.9763 |
| Space group | $P2_12_12_1$ |
| Unit cell dimensions |  |
| a,b,c (Å) | 40.5, 47.4, 151.9 |
| $\alpha,\beta,\gamma$ (°) | 90, 90, 90 |
| Total reflections | 242242 (19836) |
| Unique reflections | 122957 (11329) |
| Multiplicity | 2.0 (1.8) |
| Completeness (%) | 99 (92) |
| I/$\sigma$ | 12.89 (2.75) |
| R-meas | 0.0424 (0.293) |
| CC1/2 | 0.998 (0.902) |
| **Refinement** |  |
| Resolution range (Å) | 40.2 - 1.09 (1.13 - 1.09) |
| R-work | 0.154 (0.221) |
| R-free | 0.173 (0.216) |
| RMSD bonds (Å) | 0.01 |
| RMSD angles (°) | 1.32 |
| Wilson B-factor (Å<sup>2</sup>) | 9.65 |
| Ramachandran favored (%) | 97.8 |
| Ramachandran allowed (%) | 2.2 |

Values in parentheses indicate highest-resolution shell.

**Figure 3.**
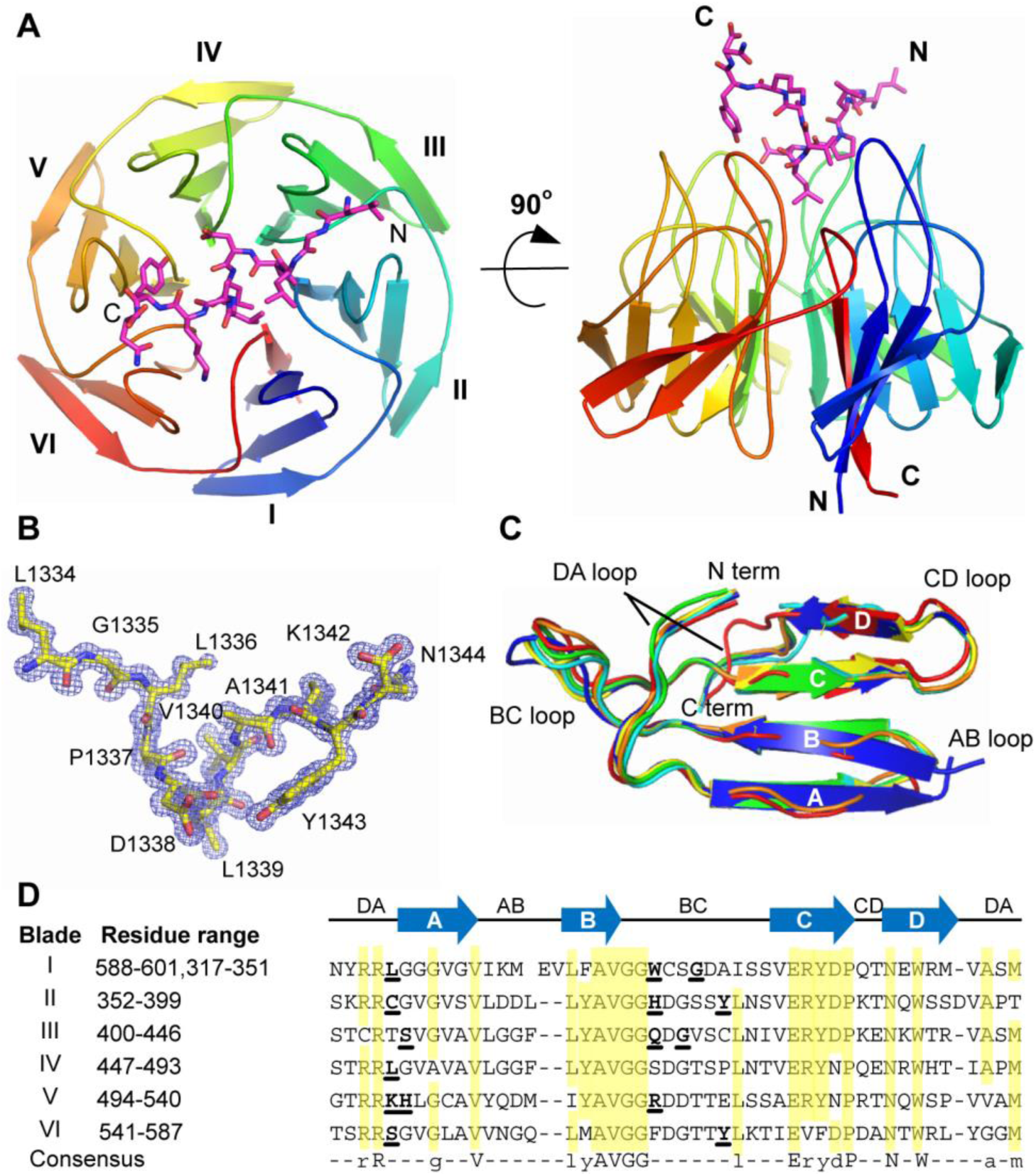
High resolution structure of KLHL20 bound to DAPK1 peptide. **(A)** Overview of the structure of KLHL20 Kelch domain in complex with DAPK1 peptide (purple sticks). Kelch repeats forming blades I to VI are labelled. **(B)** 2Fo-Fc electron density map (blue mesh) for the DAPK1 peptide contoured at 1.0 σ. **(C)** Superposition of Kelch domain blades I to VI colored from blue to red. Each blade is composed of four antiparallel β-strands (labelled A-D) and connecting loops. **(D)** Sequence alignment of the six Kelch repeats in KLHL20. Conserved residues are highlighted in yellow. DAPK1-interacting residues are shown in bold and underlined.

The Kelch domain structure shows a canonical β-propeller fold. The six Kelch repeats form the six blades (I-VI) of the propeller arranged radially around a central axis. Each repeat is folded into a twisted β-sheet consisting of four antiparallel β-strands (A-D, Figure 3C). A final C-terminal β-strand is observed to close the β-propeller and inserts into blade I as the innermost βA strand. Blade I is therefore compromised of a C-terminal βA strand and N-terminal βB, βC, and βD strands. Packing between each blade is mediated by a number of conserved hydrophobic positions as well as several buried charged residues that recur within each kelch repeat (Figure 3D).

The substrate binding surface on KLHL20 is shaped by the long BC loops, which protrude outwards from the Kelch domain surface, and the largely buried DA loops which link adjacent blades and contribute to the protein core. Notably, the six BC loops in KLHL20 are all of equal length comprising 11 residues (Figure 3D), whereas other kelch domain structures have shown more varied loop lengths across the different blades (Canning et al., 2013).

### Extended interactions of the DAPK1 peptide

The bound DAPK1 peptide shows an extended conformation that packs between Kelch domain blades II and III at its N-terminus and blades V and VI at its C-terminus (Figure 4A). At its centre, the peptide adopts a single loose helical turn that is stabilized by intramolecular hydrogen bonds between the carbonyl of Pro1337 and the amides of Val1340 and Ala1341. Here, the peptide inserts deeply into the central cavity of the Kelch domain β-propeller where it is anchored in the complex by Leu1339, the second leucine in the ‘LPDLV’ motif (Figure 4B). Binding at this central region allows the peptide to form additional contacts with blades I and IV. Thus, DAPK1 forms interactions with all six Kelch repeats, including interactions with all six DA loops and all BC loops with the exception of the BC loop in blade IV (Figures 3D and 4C).

**Figure 4.**
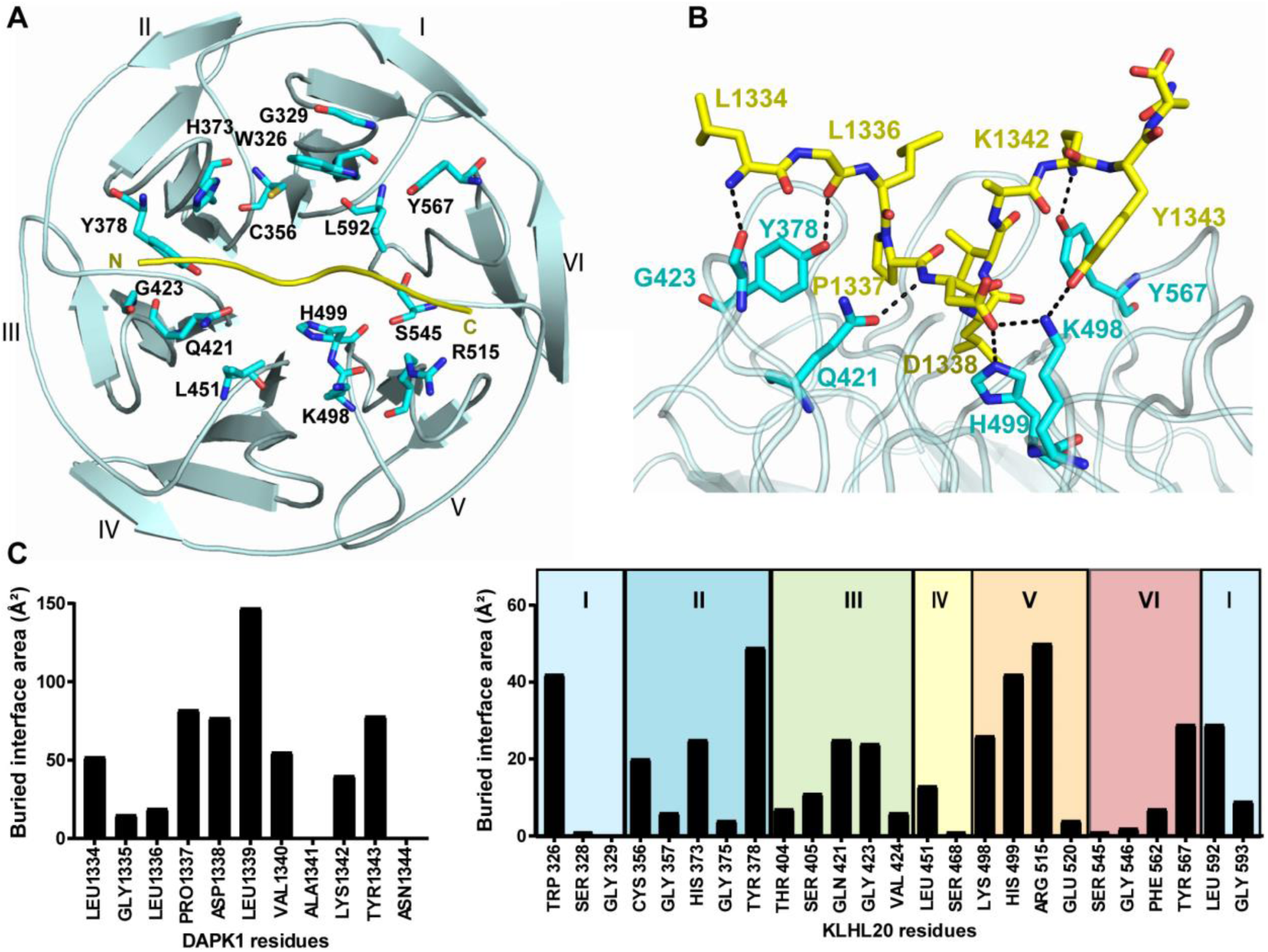
Interactions in the KLHL20-DAPK1 complex. **(A)** An overview of the DAPK1-binding residues in KLHL20 colored cyan with DAPK1 peptide shown as yellow ribbon. **(B)** Salt bridge and hydrogen bond interactions in the complex interface are shown by dashed lines. **(C)** Buried interface surface areas for interacting residues in the KLHL20-DAPK1 complex.

### Interactions of the ‘LPDLV’ motif

The ‘LPDLV’ motif of DAPK lies at the core of the protein-peptide interface. Here, the hydrophobic side chains pack against Kelch domain blades I and II and make notable van der Waals contacts with KLHL20 Trp326, His373 and Leu592, respectively (Figure 5A). Somewhat surprisingly the first leucine in the ‘LPDLV’ motif, Leu1336, is oriented away from the binding interface and has only minor interaction with KLHL20, mostly through its main chain atoms. In the SPOT peptide arrays, changes at this position reduced KLHL20 binding significantly, but did not abolish it (Figure 2). The importance of this position likely stems from the conformational constraints of the following DAPK1 residue Pro1337. By contrast, the second leucine, Leu1339, is the most buried DAPK1 residue in the complex (Figure 4C). This side chain lies sandwiched between KLHL20 His499 and Leu592 (Figures 5A-B), but forms interactions across all the Kelch repeats, except for blade IV, by virtue of its central binding position. The final residue in the ‘LPDLV’ motif, DAPK1 Asp1338, is oriented away from the hydrophobic side chains to face Kelch domain blade V, where it forms a salt bridge with KLHL20 Lys498, as well as a hydrogen bond to His499 (Figure 5B). Residues across the mapped DAPK1 binding motif are well conserved across species (Figure 5C).

**Figure 5.**
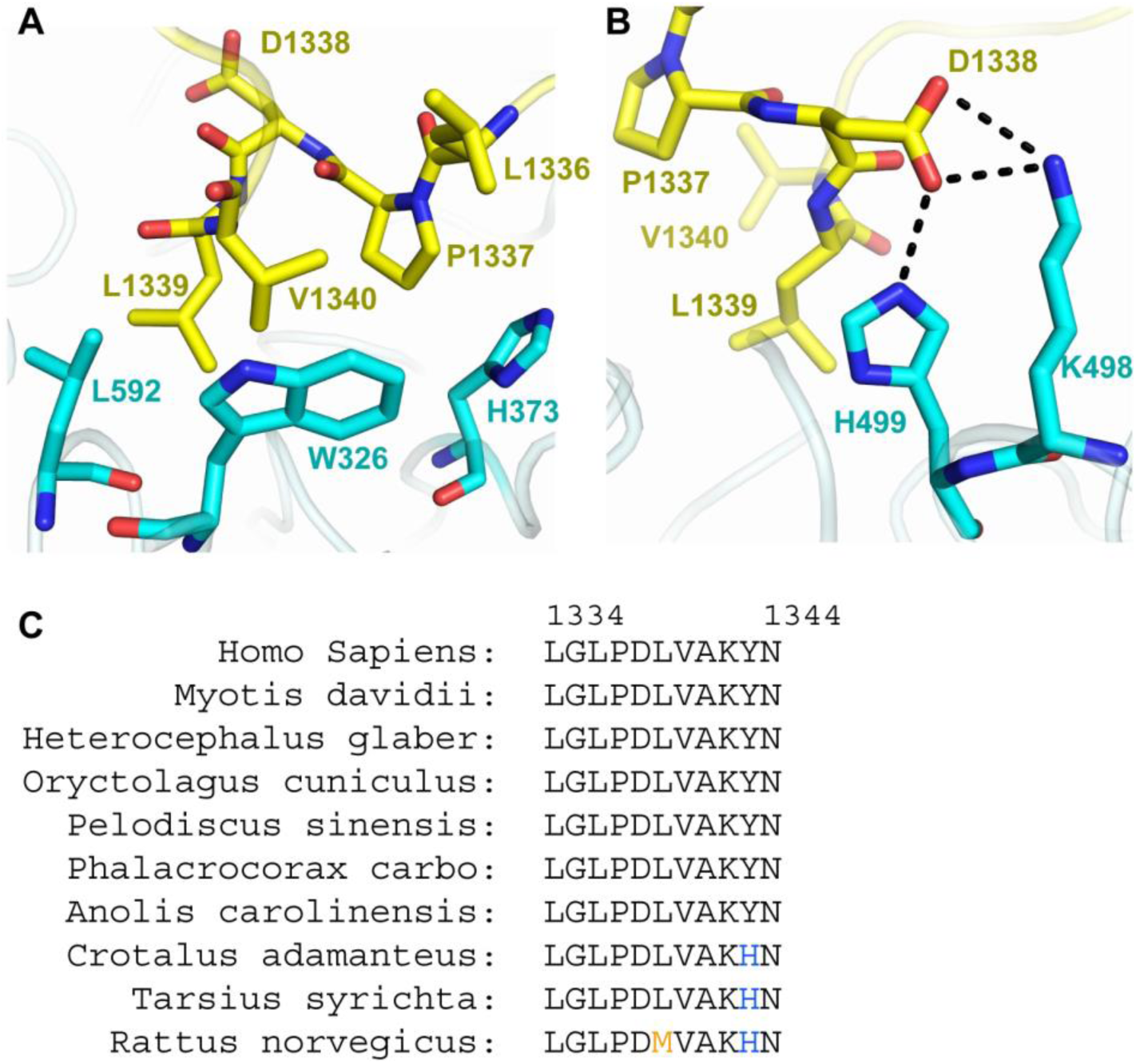
Interactions of the DAPK1 ‘LPDLV’ motif. (**A**) Hydrophobic interactions between the DAPK1 ‘LPDLV’ motif and KLHL20 Kelch domain blades I and II. (B) DAPK1 Asp1338 forms a salt bridge with KLHL20 Lys498, as well as a hydrogen bond to H499. (**C**) The sequence of the crystallized DAPK1 peptide is conserved across species.

### Direct and water-mediated hydrogen bonding in the KLHL20-DAPK1 complex

In total, the complex between KLHL20 and DAPK1 includes 8 direct hydrogen bond or salt bridge interactions (Figure 4B), as well as a number of water-mediated interactions (Figure 6). The N-terminal three residues of the DAPK1 peptide are oriented away from the KLHL20 surface. Their binding interactions are mediated by their main chain atoms, which form hydrogen bonds with KLHL20 residues Tyr378, Gln421 and Gly423, respectively (Figure 4B). Gln421 forms an additional hydrogen bond with the backbone amide of DAPK1 Asp1338, one of the critical residues within the ‘LPDLV’ motif. Towards the C-terminus of the peptide, interactions are formed through DAPK1 Lys1342 and Tyr1343, while Ala1341 and Asn1344 are oriented to solvent. Lys1342 folds towards Kelch domain blade VI where it forms a direct hydrogen bond to the BC loop residue Tyr567 (Figure 4B). DAPK1 Tyr1343 folds instead against blade V to hydrogen bond with KLHL20 Lys498 (DA loop, Figure 4B) and forms additional hydrophobic packing with the BC loop residue Arg515. Water molecules in the complex help to bridge more distant contacts or to satisfy other nitrogen and oxygen atoms that otherwise lack direct hydrogen bonds (Figure 6).

**Figure 6.**
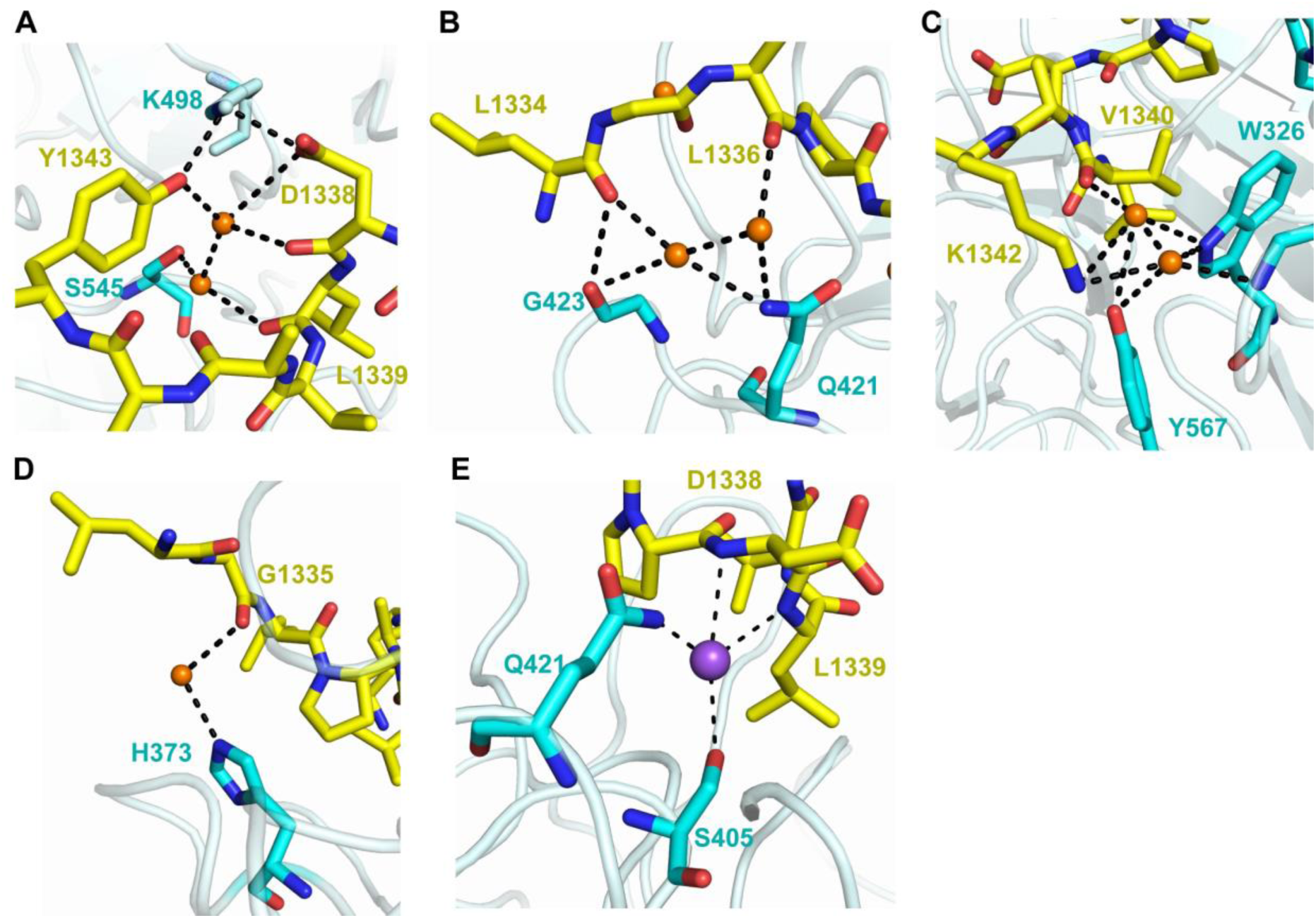
Indirect contacts between KLHL20 and DAPK1. **(A-D)** Water-bridged hydrogen bonds in the protein-peptide interface. Waters are shown as orange spheres. **(E)** A sodium ion (purple sphere) is proximal for electrostatic interactions with the side chains of KLHL20 Ser405 and Gln421 as well as the backbone atoms of DAPK1 D1338 and L1339.

## DISCUSSION

The Cullin-RING E3 ligase KLHL20 has been shown to ubiquitinate some half a dozen protein targets that link its activities to diverse processes including autophagy, hypoxia, cancer and Alzheimer’s disease (Chen et al., 2016). Here, we performed the first structural and biochemical analyses of KLHL20 to elucidate how it engages its substrates through the Kelch β-propeller domain. Structural studies required the identification of a short DAPK1 peptide motif that subsequently enabled crystallization. As a result, we were able to solve the structure of the KLHL20-DAPK1 complex at 1.1 Å resolution. The structure identifies a central ‘LPDLV’ motif in the DAPK1 epitope that inserts into the central pocket of the Kelch β-propeller as a loose helical turn. The interface in KLHL20 complements this motif with a hydrophobic core supported by a salt bridge interaction. The recognition motifs within other KLHL20 substrates remain to be defined at the same level, but are likely to form a similar pattern of hydrophobic and charge-charge interactions.

The low micromolar binding of the DAPK1 peptide to KLHL20 is comparable in affinity to substrates of the SPOP E3 ligase (Zhuang et al., 2009), which similarly assembles into a Cullin-RING ligase complex through CUL3 (Errington et al., 2012). However, this is weaker than the low nanomolar binding observed for NRF2 interaction with the Kelch domain of KEAP1 (Tong et al., 2006). These differences may reflect the strict regulation of NRF2, which ensures its constitutive degradation. Alternatively, there may be differences in affinity between the binding of DAPK peptide and the death domain in the context of the full length DAPK1 protein. Death domains are well known protein interaction modules that fold as a bundle of six α-helices (Ferrao and Wu, 2012). While the isolated death domain of DAPK1 appears to be intrinsically disordered, it is possible that other DAPK regions contribute to its proper folding (Dioletis et al., 2013). The critical ‘LPDLV’ motif of DAPK1 maps to the predicted α3 helix. To understand how this might interact with KLHL20 in the context of the full death domain we built a homology model of human DAPK1 using the MyD88 protein structure (PDB 3MOP) as a template and ICM-Pro software (Molsoft) (Figures 7A-B) (Abagyan et al., 1997). Superposition of the ‘LPDLV’ motifs revealed good agreement between the model and our crystal structure (Figures 7B-C). Overall, the helical turn of the DAPK1 peptide was a good match to the folding of the α3 helix (Figures 7C-7D). Consequently, the side chains in the respective structures were closely aligned (Figure 7C). Importantly, the key interacting residues of DAPK1 were exposed providing a surface epitope for KLHL20 to bind. The model suggests that the α3 helix of the death domain can insert into the relatively wide pocket of KLHL20 to recapitulate the observed peptide interaction without steric hindrance. There is some precedent for such an arrangement from the structure of KEAP1 bound to the DLG motif of NRF2, which also formed an extended helical structure (Fukutomi et al., 2014).

**Figure 7.**
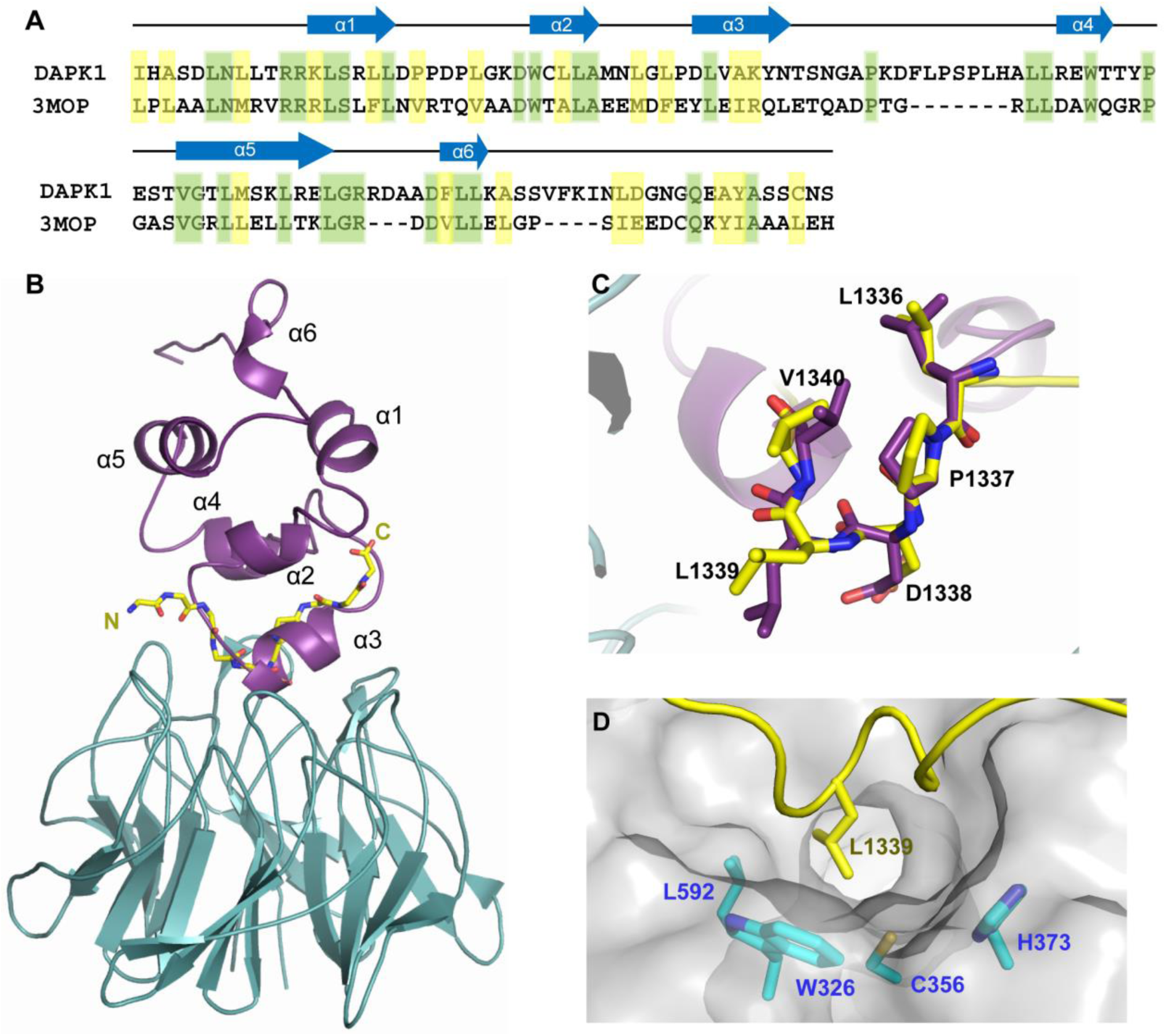
Homology model of DAPK1 death domain in complex with KLHL20. **(A)** Sequence alignment of the death domains of DAPK1 and MyD88 (PDB 3MOP, chain A). **(B)** Superposition of the KLHL20-DAPK1 structure (cyan/yellow) and a homology model of the DAPK1 death domain (purple; template PDB 3MOP) based on the critical ‘LPDLV’ motifs. **(C)** Close-up view showing good agreement between the helical conformations of the ‘LPDLV’ motif in the crystallized DAPK1 peptide and the homology model (α3). **(D)** Surface representation of KLHL20 highlighting the potentially druggable pocket bound by DAPK1 Leu1338. In addition to hydrophobics such as KLHL20 Trp326, the proximal location of KLHL20 Cys356 suggests opportunity for the development of covalent inhibitors.

Of note, previous studies of KEAP1 have characterized the binding of both unmodified and phosphorylated peptides (for example, the ‘ETGE’-containing motif from NRF2 and the ‘phospho-STGE’-containing motif of sequestosome-1/p62)(Ichimura et al., 2013; Lo et al., 2006). To date, no posttranslational modifications have been reported for the death domain of DAPK1 (www.phosphosite.org; (Hornbeck et al., 2015)) and we have yet to identify a phosphorylated substrate motif for KLHL20. Nonetheless, other substrates of KLHL20 may similarly substitute a phosphorylated residue for the aspartate found in the ‘LPDLV’ motif of DAPK1. It is known for example that KLHL20 binds specifically to the activated pool of ULK1 to terminate autophagy (Liu et al., 2016).

KLHL20 has emerged as an interesting target for drug development with potential application in both oncology and Alzheimer’s disease. Inhibition of KLHL20 would help to stabilize the tumor suppressor proteins DAPK1 and PML (Lee et al., 2010; Yuan et al., 2011). It could also stabilize ULK1 to prolong autophagy allowing greater clearance of potentially toxic misfolded proteins (Liu et al., 2016). The structure of KLHL20 at atomic resolution provides a strong template for structure-based drug design. Moreover, the identified DAPK1 peptide provides a valuable reagent for drug screening assays based on peptide displacement. Of note, the hydrophobic interaction surface in KLHL20 includes an exposed cysteine residue (Cys356) that lies within 4 Å of the bound peptide (Figure 7D). This cysteine is accessible for modification and proximal to KLHL20 Trp326, another key DAPK1 interacting residue (Figures 5A and 7D). Thus, KLHL20 may be a promising target for screening against covalent inhibitor or fragment libraries.

To date, few Kelch-substrate complexes have been structurally characterized, with the major examples being the KEAP1-NRF2 (Fukutomi et al., 2014; Lo et al., 2006) and KLHL3-WNK4 systems (Schumacher et al., 2014). The Kelch family proteins are relatively diverse in their primary sequences. Indeed, KLHL20 shares only 25 to 50% sequence identity with other human Kelch domains. Comparison of the available complex structures shows that the substrate binding surfaces in KEAP1 and KLHL3 are largely polarized to opposite sides of the β-propeller, whereas the KLHL20 interface is more central (Figure 8A). The polarized binding is supported by variability in the BC loop lengths, which is not found in KLHL20. The patterning of hydrophobic and charged residues also differs across the different structures (Figures 8B-D). The interaction surface in KLHL20 provides a more substantial hydrophobic contribution that should make it more favorable for inhibitor development.

**Figure 8.**
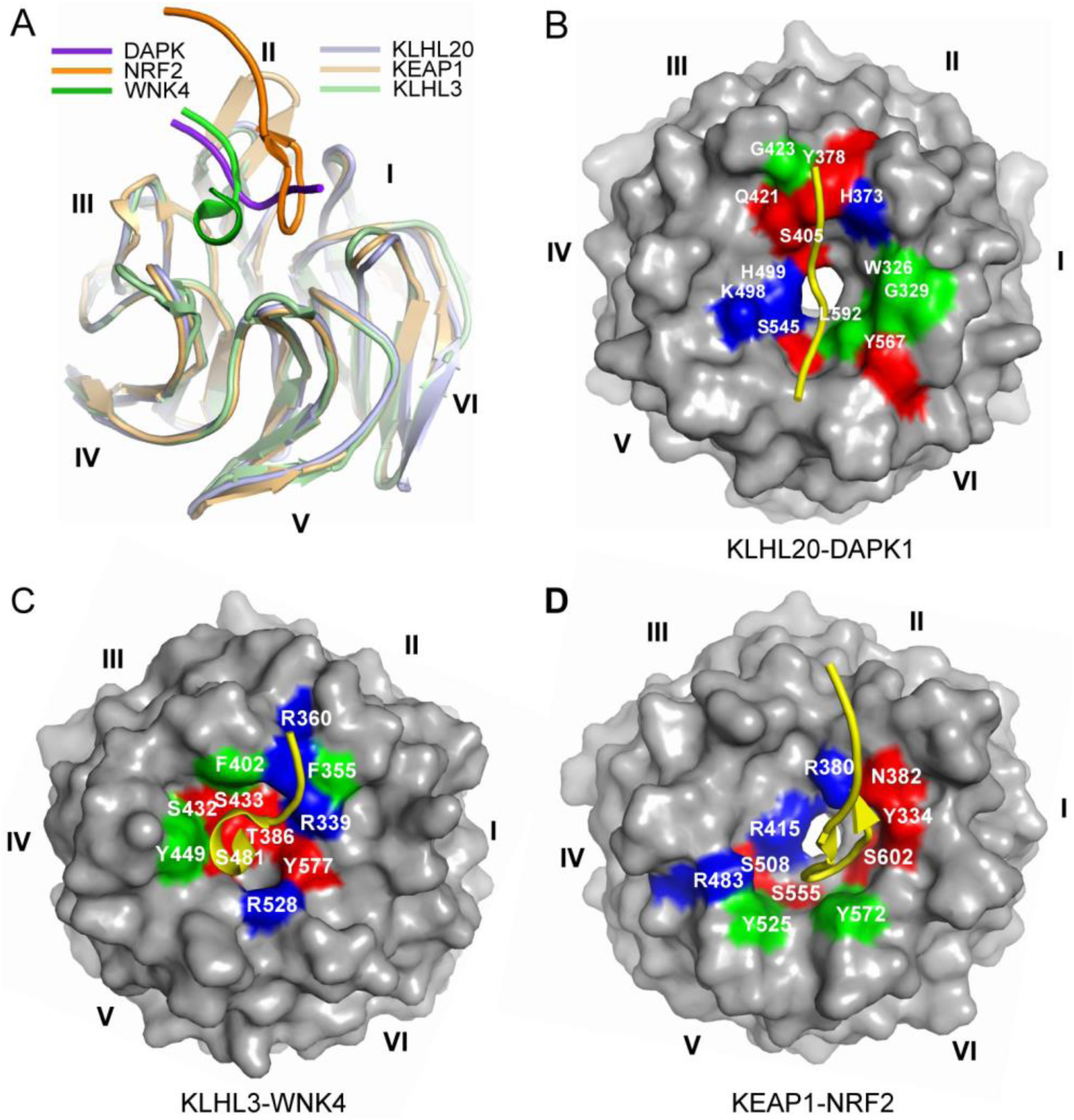
The binding mode of DAPK1 is distinct from other Kelch-substrate complexes. (**A**) Superposition of the complex structures of KLHL20-DAPK1 (light and dark purple), KEAP1-NRF2 (PDB 2FLU, light and dark orange) and KLHL3-WNK4 (PDB 4CH9, light and dark green). (**B**-**D**) Surface representations of the Kelch domains in the KLHL20-DAPK1 (**B**), KLHL3-WNK4 (**C**) and KEAP1-NRF2 (**D**) complexes with key contact residues highlighted by their binding characteristics (blue, basic residue; red, other polar; green, hydrophobic). The distinct surfaces and bound peptide conformations (yellow ribbons) highlight the rich variety of binding modes that can be established by the circular Kelch domain substrate pockets.

For the most part, the study of protein-peptide complexes has focused on protein interaction domains with linear binding grooves. The Kelch β-propeller provides an example of a circular pocket that can accommodate peptides with unexpected turns, twists and helices. Overall, this feature is likely to increase the diversity of substrate interaction modes and allow for more selective drug design. This is perhaps reflected in the prevalence of Kelch and WD40 domains within the family of Cullin-RING E3 ligases.

## ACKNOWLEDGEMENTS

The authors would like to thank Diamond Light Source for beamtime (proposal mx15433), as well as the staff of beamline I03 and I24 for assistance with crystal testing and data collection. The SGC is a registered charity (number 1097737) that receives funds from AbbVie, Bayer Pharma AG, Boehringer Ingelheim, Canada Foundation for Innovation, Eshelman Institute for Innovation, Genome Canada, Innovative Medicines Initiative (EU/EFPIA) [ULTRA-DD grant no. 115766], Janssen, Merck KGaA Darmstadt Germany, MSD, Novartis Pharma AG, Ontario Ministry of Economic Development and Innovation, Pfizer, São Paulo Research Foundation-FAPESP, Takeda, and Wellcome [106169/ZZ14/Z].

## AUTHOR CONTRIBUTIONS

A.N.B. and V.D’A. designed and supervised the research. S.P. and P.F. printed the peptide arrays. Z.C. performed the experiments and solved the crystal structure. Z.C. and A.N.B wrote the paper.

## STAR Methods

### KEY RESOURCES TABLE

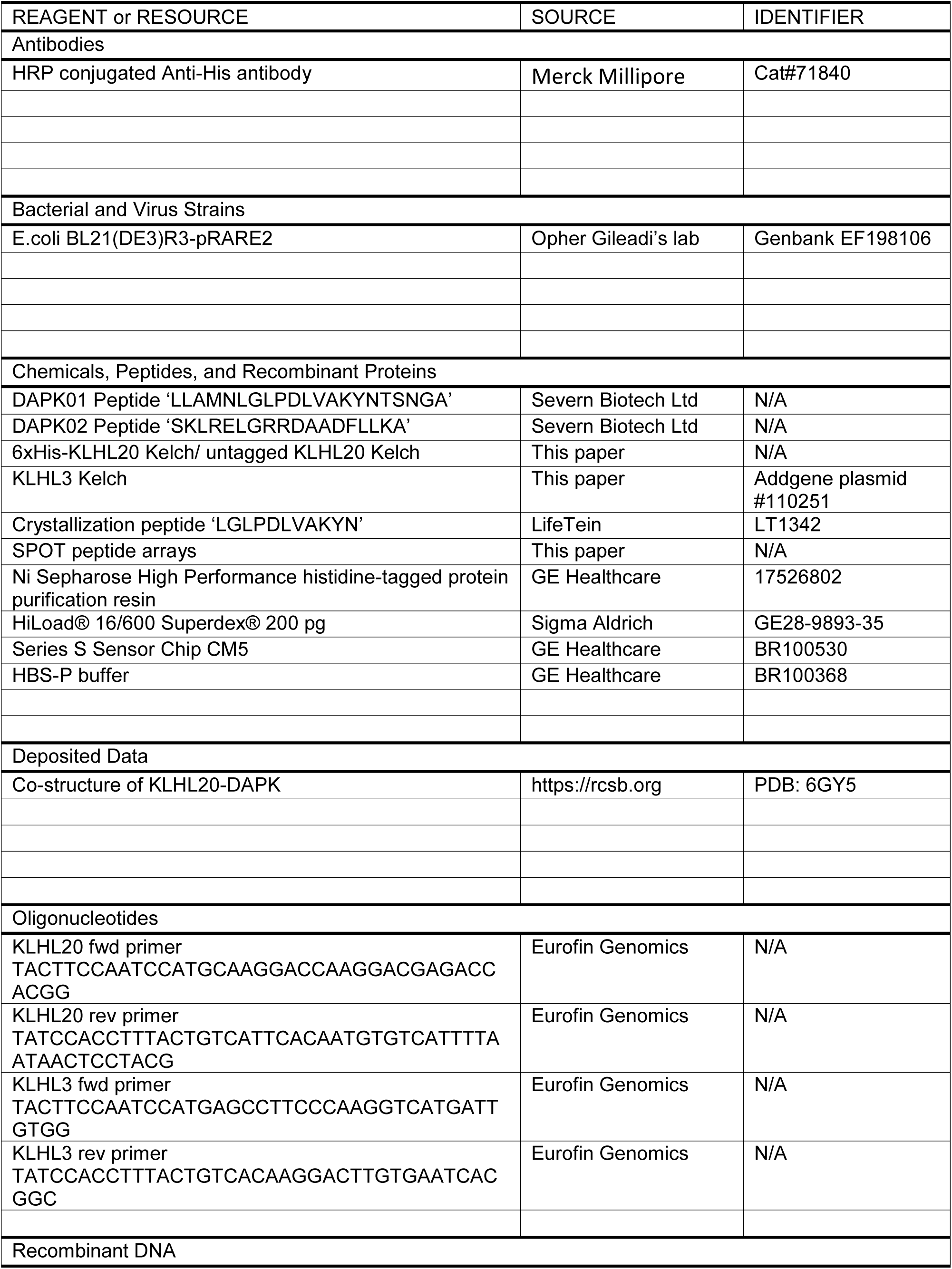
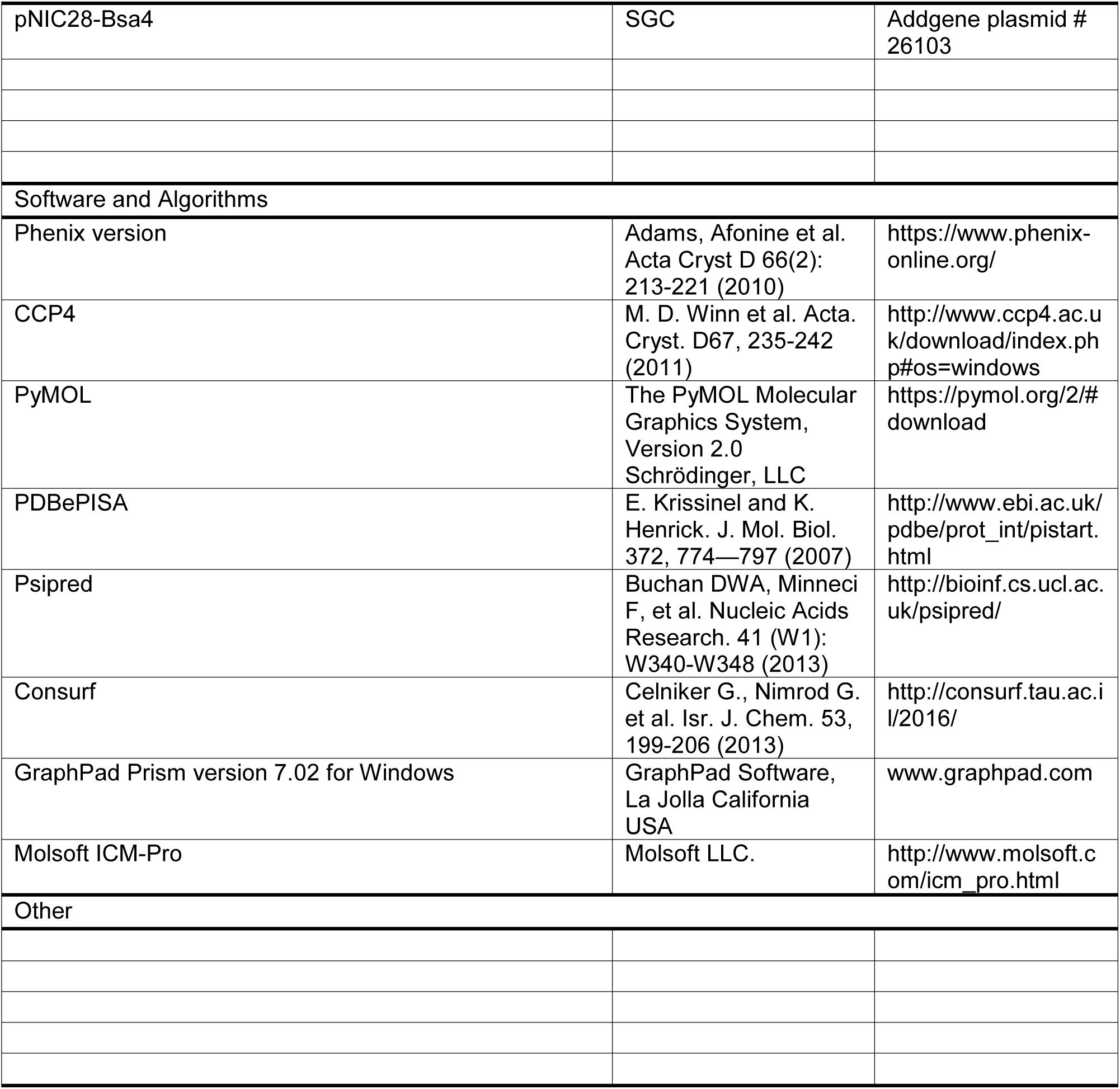

### CONTACT FOR REAGENT AND RESOURCE SHARING

Further information and requests for resources and reagents should be directed to and will be fulfilled by the Lead Contact, Alex Bullock

## METHOD DETAILS

### Constructs

The Kelch domains of human KLHL20 (Uniprot Q9Y2M5 isoform 1, M303-E605) and KLHL3 (Uniprot Q9UH77 isoform 1, residues S298–L587) were cloned using ligation-independent cloning into the bacterial expression vector pNIC28-Bsa4 (GenBank accession number EF198106), which provides for an N-terminal hexahistidine tag and TEV cleavage site. DNA sequences were verified by Source Bioscience Ltd.

### Protein expression and purification

Plasmids were transformed into *E. coli* strain BL21(DE3)R3-pRARE2. Cells were cultured in LB broth at 37°C until OD_600_ reached 0.6. Recombinant protein expression was then induced by addition of 0.4 M isopropyl β-D-1-thiogalactopyranoside, followed by 18 hours continuous shaking at 18°C. Cells were harvested by centrifugation and lysed by sonication in binding buffer (50 mM HEPES pH 7.5, 500 mM NaCl, 5% glycerol, 5 mM imidazole) supplemented with 0.5 mM TCEP. Recombinant proteins were captured on nickel sepharose resin, washed with binding buffer and eluted by a stepwise gradient of 30-250 mM imidazole. Further clean-up was performed by size exclusion chromatography using a HiLoad 16/60 S200 Superdex column buffered in 50 mM HEPES pH 7.5, 300 mM NaCl, 0.5 mM TCEP. Finally, the eluted protein was purified by anion exchange chromatography using a 5 mL HiTrap Q column. Protein masses were confirmed by intact LC-MS mass spectrometry. Where required, the hexahistidine tag was cleaved overnight at 4°C using TEV protease.

### Peptide arrays (SPOT assay)

Cellulose-bound peptide arrays were prepared employing standard Fmoc solid phase peptide synthesis using a MultiPep-RSi-Spotter (INTAVIS, Köln, Germany) as previously described (Picaud and Filippakopoulos, 2015). After array synthesis, membranes were incubated with 5% BSA to block free sites. The arrays were then incubated with 1 μM recombinant hexahistidine-tagged KLHL20 Kelch domain in PBS at 4°C overnight. Unbound protein was washed off in PBS buffer with 0.1% Tween 20 and bound protein was detected using HRP-conjugate anti-His antibody.

### Surface Plasmon Resonance

Assays were performed at 25°C using a BIACORE S200 (GE Healthcare) surface plasmon resonance (SPR) instrument. The Kelch domains of KLHL20 and KLHL3 were immobilized on sensor chip CM5 (GE Healthcare) using amine coupling. Reference flow cells had no immobilized protein. Binding was monitored using a flow rate of 30 μL/min. The peptide analytes were prepared in HBS-P buffer (GE Healthcare). Data reported were after reference flow cell signal subtraction. Data were analyzed by one-site steady-state affinity analysis using the Biacore S200 Evaluation software and the fitting equation 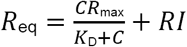. (RI, bulk refractive index contribution; *K*_D_, dissociation constant; C, analyte concentration).

### Protein crystallization

The purified KLHL20 protein was concentrated to 12 mg/mL using a 10 kDa molecular-mass cut-off centrifugal concentrator in 50 mM HEPES pH 7.5, 300 mM NaCl and 5 mM TCEP buffer. The 11-residue DAPK1 peptide (LGLPDLVAKYN) was added in the same buffer to a final concentration of 3 mM. The protein-peptide mixture was incubated on ice for 1 hour prior to setting up sitting-drop vapour-diffusion crystallization plates. Micro-seed stocks were prepared from small KLHL20 crystals grown during previous rounds of crystal optimization. Those early crystals were transferred into an Eppendorf tube containing 50 μL reservoir solution and a seed bead (Hampton Research), then vortexed for 2 min. Seed stocks were diluted 500 fold before use. The best-diffracting crystals of the KLHL20 complex were obtained at 20°C by mixing 75 nL protein, 20 nL diluted seed stock and 75 nL of a reservoir solution containing 2 M sodium chloride and 0.1 M acetate buffer pH 4.5. Prior to vitrification in liquid nitrogen, crystals were cryoprotected by direct addition of reservoir solution supplemented with 25 % ethylene glycol.

### Structure determination

Diffraction data for the KLHL20-DAPK1 complex were collected on beamline I03 at Diamond Light Source, Didcot, U.K. Data were processed in PHENIX version1.9 (Adams et al., 2010). Molecular replacement was performed with PHENIX.Phaser-MR using KLHL12 (PDB 2VPJ chain A) as the search model. PHENIX.Autobuild was used to build the initial structural model. COOT (Emsley et al., 2010) was used for manual model building and refinement, whereas PHENIX.REFINE was used for automated refinement. TLS parameters were included at later stages of refinement. Tools in COOT, PHENIX and MolProbity (Chen et al., 2010) were used to validate the structures.

### Homology model

A homology model for the death domain of human DAPK1 was built in Molsoft ICM-Pro software using MyD88 (PDB 3MOP chain A, 25% sequence identity) as the structural template. The initial model was refined by energy minimization and side chain optimization in ICM-Pro (Molsoft) (Abagyan et al., 1997).

